## Supplementary information for "Environmental response in gene expression and DNA methylation reveals factors influencing the adaptive potential of *Arabidopsis lyrata*"

[3] Current address: School of Natural Sciences, Massey University, Palmerston North 4442, New Zealand

[4] Current address: Embryology Research Unit, Bioinformatics Group, Children's Medical Research Institute, University of Sydney, Westmead NSW 2145, Australia

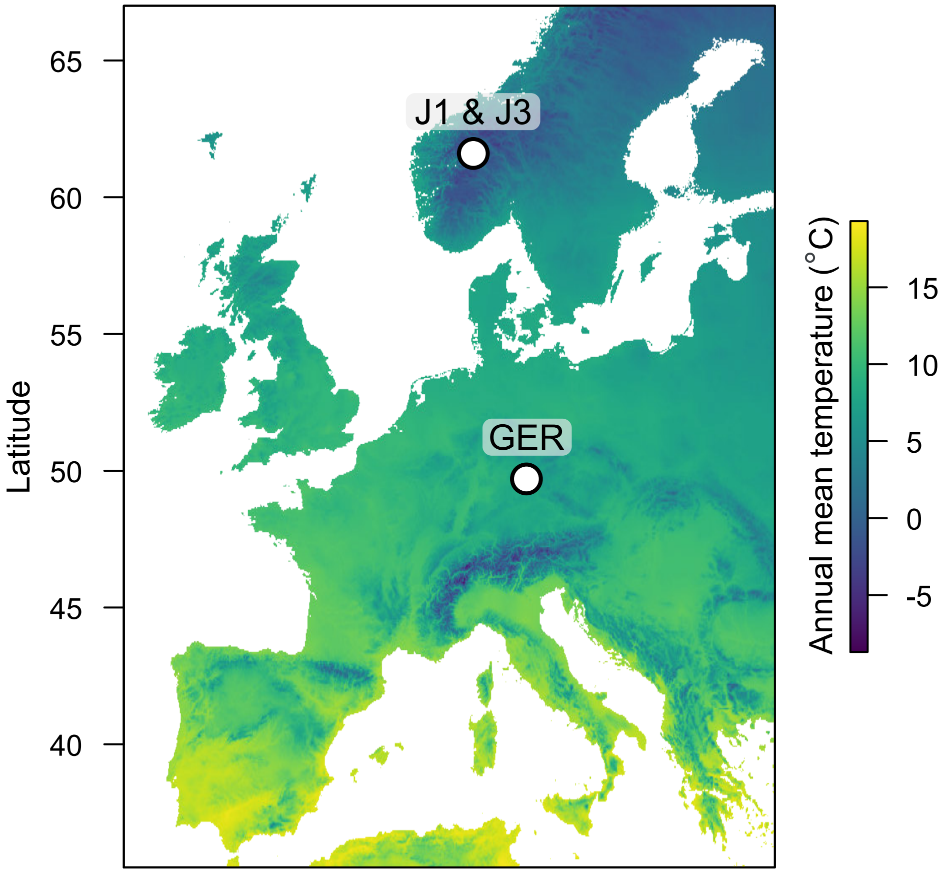


**Figure S1**. Locations of the *A. lyrata* populations. Temperature data from WorldClim.

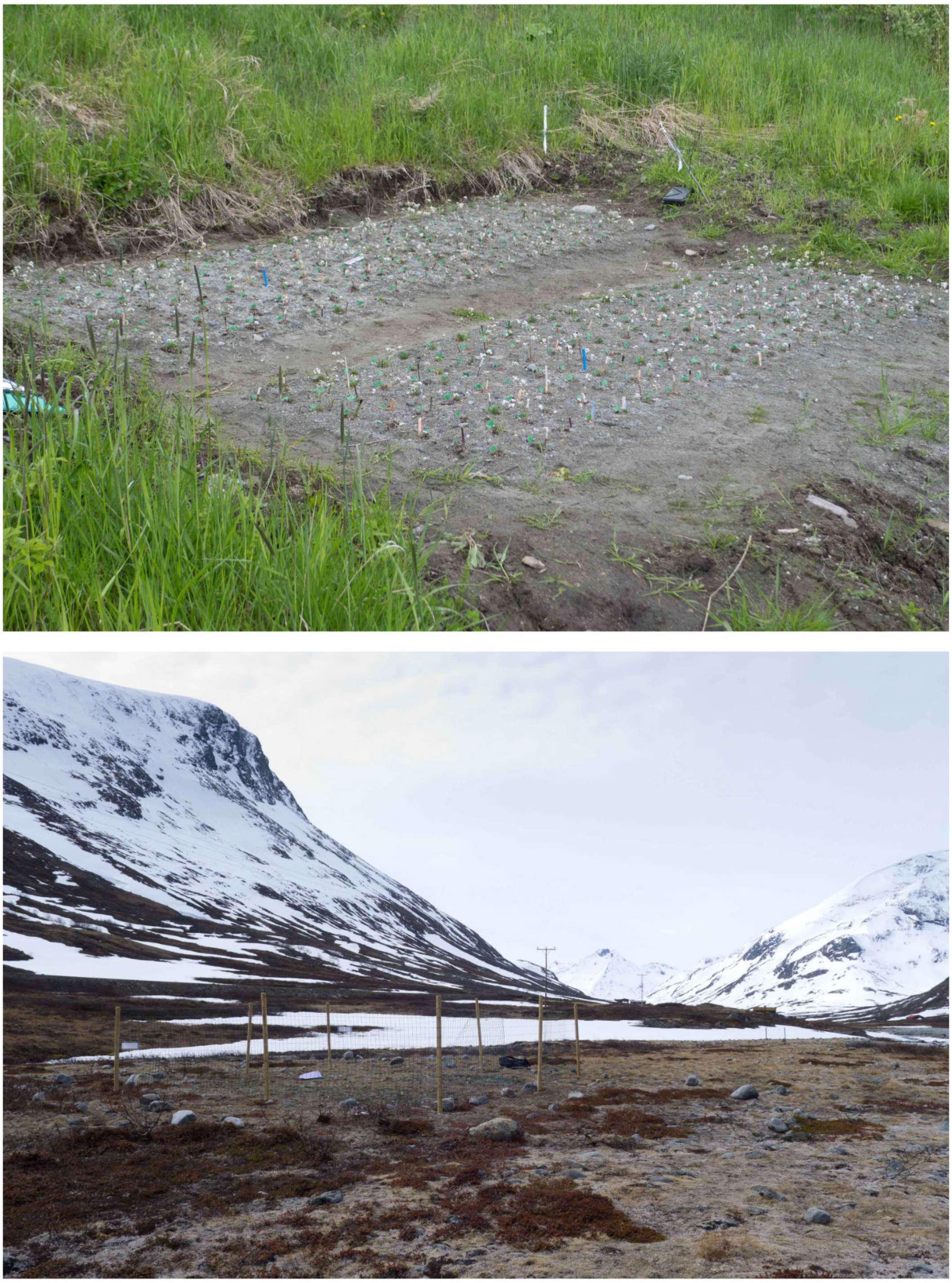

**Figure S2.** Photos of the low- and high-altitude field sites, taken during the 6$^{th}$ and 7$^{th}$ of June 2015, respectively.

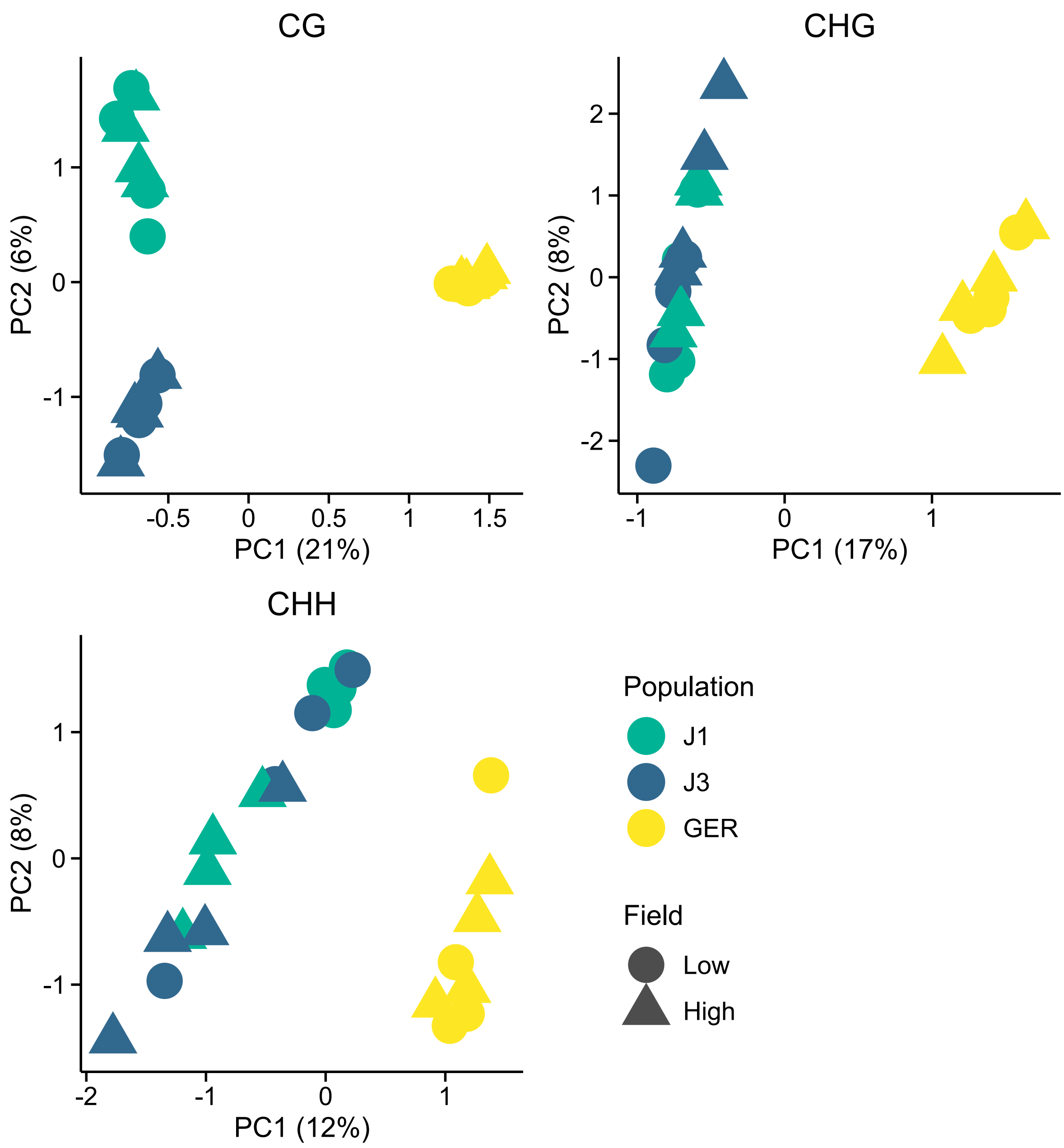


**Figure S3.** Methylation variation along the first two eigenvectors of a PCA, shown for the three methylation contexts. The proportion of variance explained by the PCs is shown in parentheses.

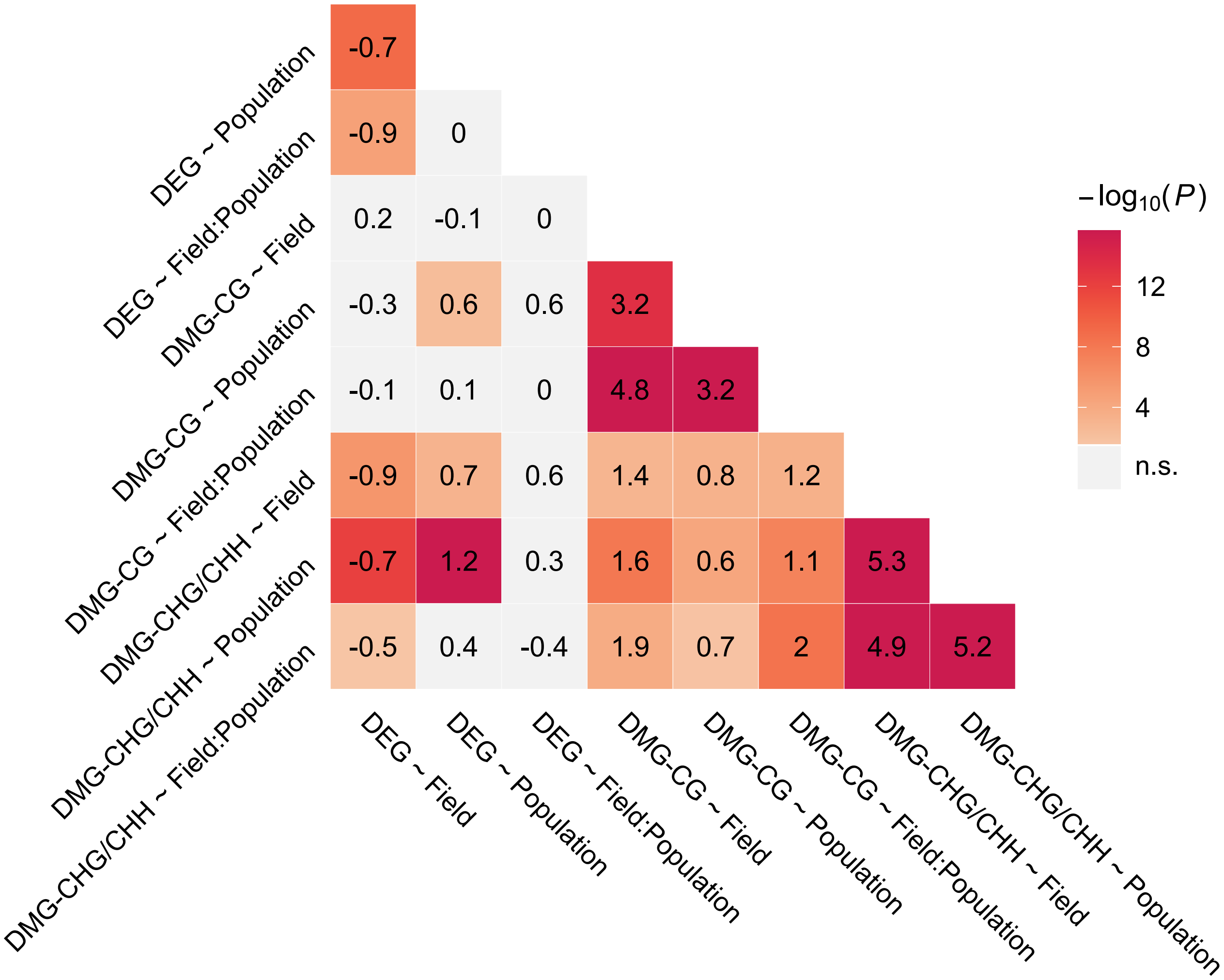


**Figure S4.** Overlap between candidate gene groups. Shown are $\log_2$ odds ratios (numbers) and *P* values (color gradient) from Fisher's exact tests.

**DEG ~ Field**

GO:0015979 Photosynthesis
GO:0015995 Chlorophyll biosynthetic process
GO:0019253 Reductive pentose-phosphate cycle
GO:0009409 Response to cold
GO:0009642 Response to light intensity

**DEG ~ Population**

GO:0042742 Defense response to bacterium
GO:0071456 Cellular response to hypoxia
GO:0051607 Defense response to virus

**DEG ~ Field:Population**

GO:0007018 Microtubule-based movement
GO:0006979 Response to oxidative stress
GO:0009407 Toxin catabolic process
GO:0071456 Cellular response to hypoxia
GO:0009409 Response to cold

**DMG-CG ~ Field**

**DMG-CG ~ Population**

GO:0031348 Neg. reg. of defense response
GO:0009833 Plant-type primary cell wall biog.
GO:0030244 Cellulose biosynthetic process

**DMG-CG ~ Field:Population**

GO:0007018 Microtuble-based movement

**DMG-CHG/CHH ~ Field**

GO:0007165 Signal transduction
GO:0006812 Cation transport
GO:0006952 Defense response

**DMG-CHG/CHH ~ Population**

GO:0007165 Signal transduction
GO:0019722 Calcium-mediated signaling
GO:0098656 Anion transmembrane transport
GO:0048544 Recognition of pollen
GO:0071704 Organic substance metabolic proc.

**DMG-CHG/CHH ~ Field:Population**

GO:0007165 Signal transduction
GO:0015969 Ammonium transport
GO:0019575 Ammonia assimilation cycle
GO:0072488 Ammonia transmembrane transport

**Figure S5**. Top five enriched ($Q < 0.05$, hypergeometric test) GO terms among each candidate gene set

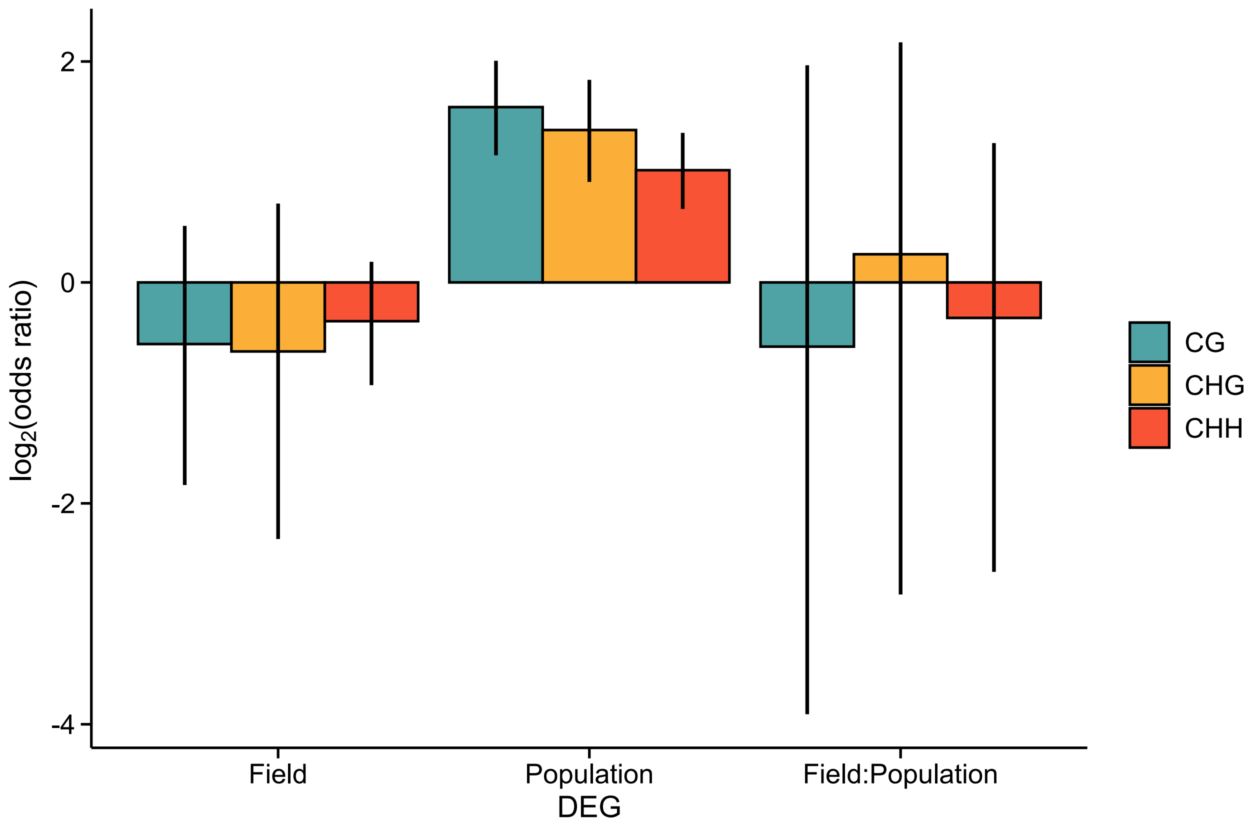


**Figure S6.** The $\log_2$ odds ratio of association between DEGs and differentially methylated regions 1 kb upstream of each gene, shown for each methylation context. Error bars show 95% CIs.

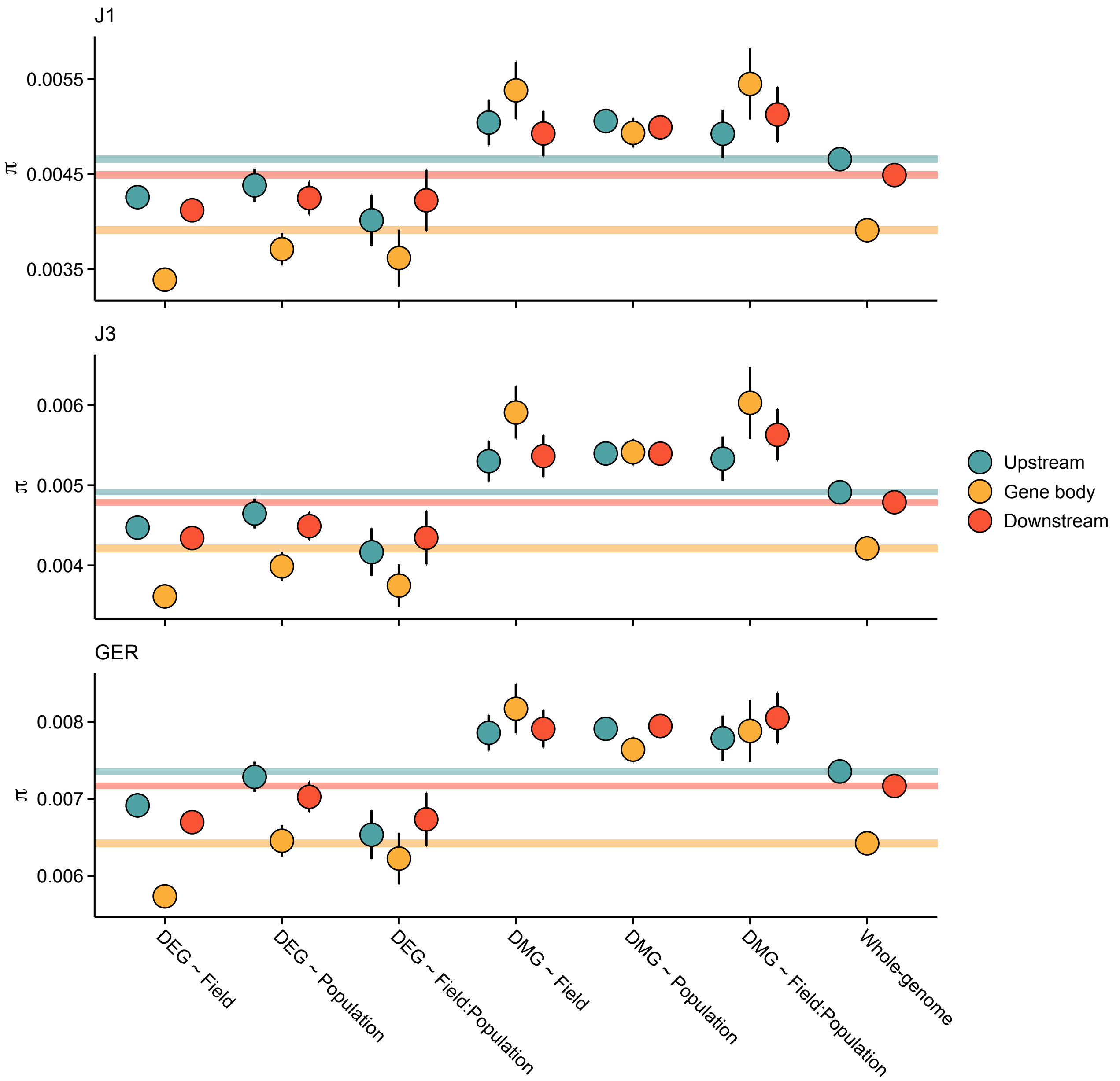


**Figure S7.** Pairwise nucleotide diversity (π) at the candidate gene sets. Shown are estimates for gene bodies and 1 kb up- and downstream regions. Error bars show 95% CIs. Shaded areas mark the 95% CIs across all genes.

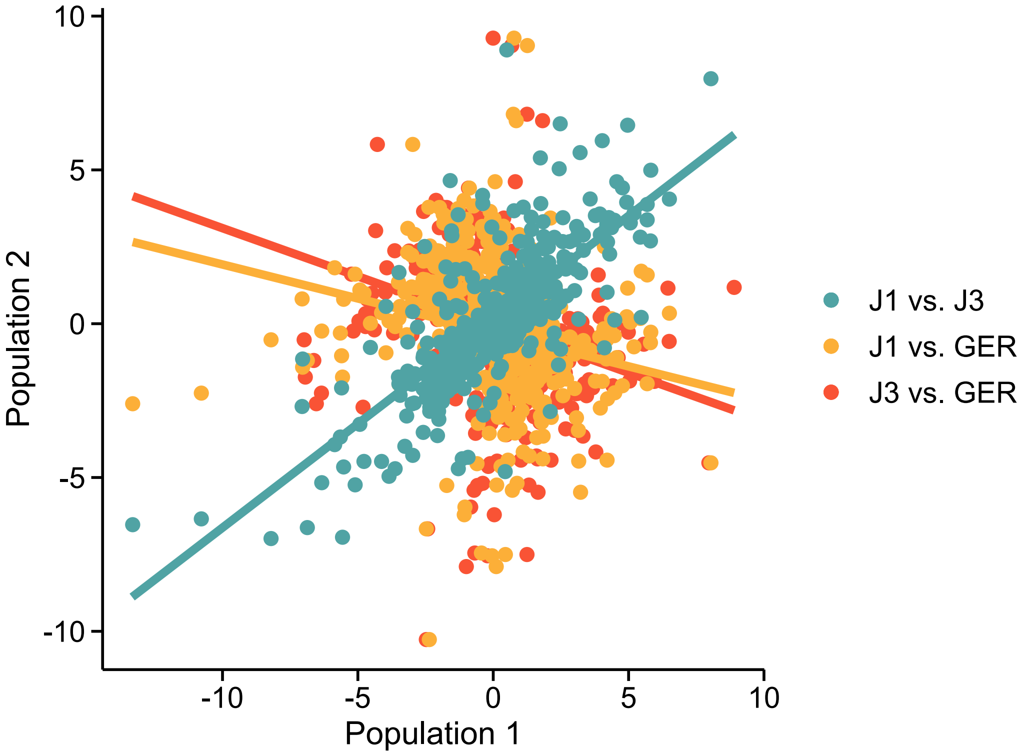


**Figure S8**. Population effects at DEG ~ Field:Population genes. For each population comparison (shown with colors), values indicate expression $\log_2$ fold changes between the low- and high-altitude field sites. For example, teal circles indicate that the fold change for J1 is on the *x*-axis and J3 on the *y*-axis. Fit from linear models are shown with lines.

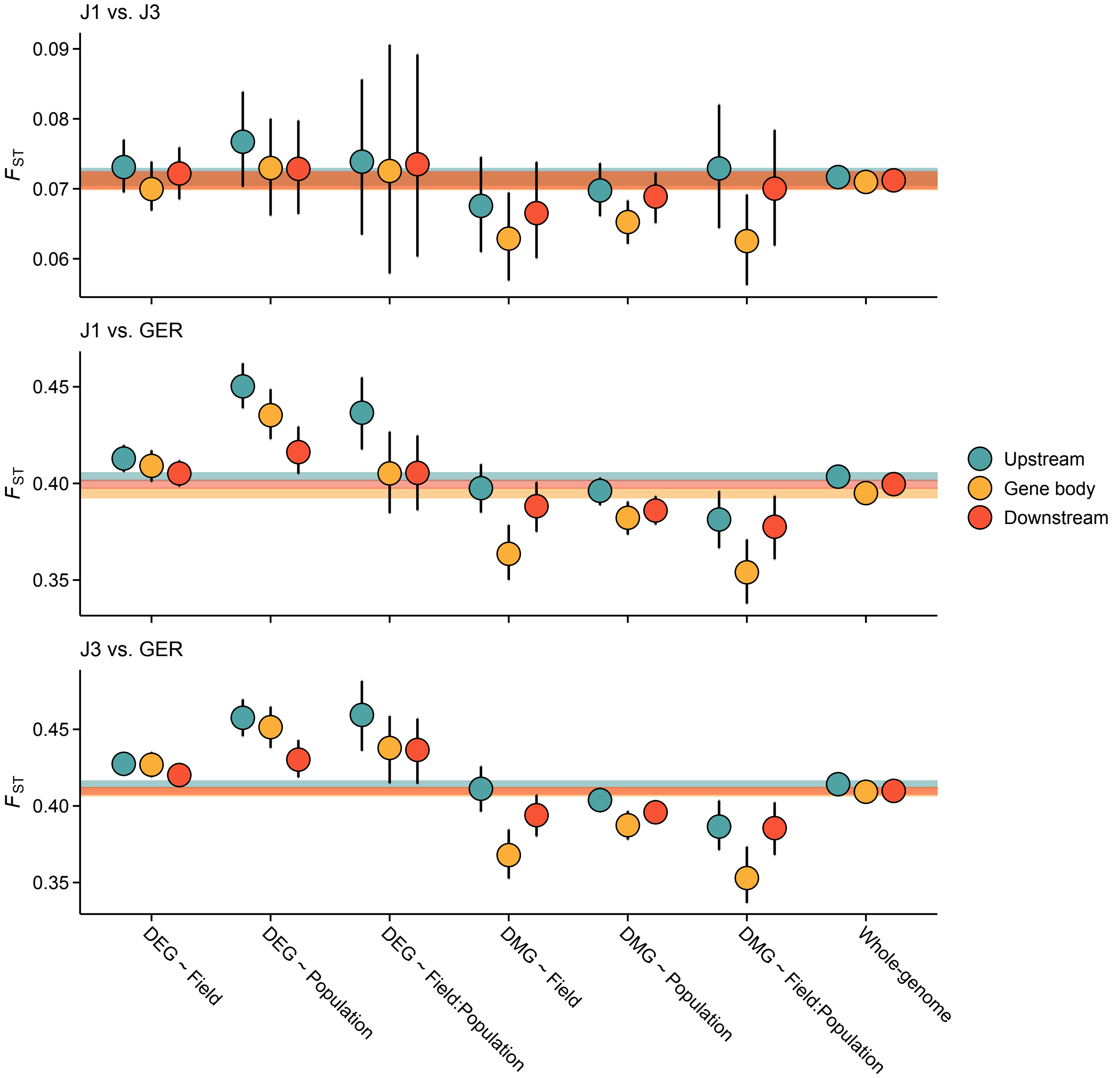


**Figure S9.** Pairwise $F_{ST}$ estimates for the candidate gene sets. Shown are estimates for gene bodies and 1 kb up- and downstream regions. Error bars show 95% CIs. Shaded areas mark the 95% CIs across all genes.

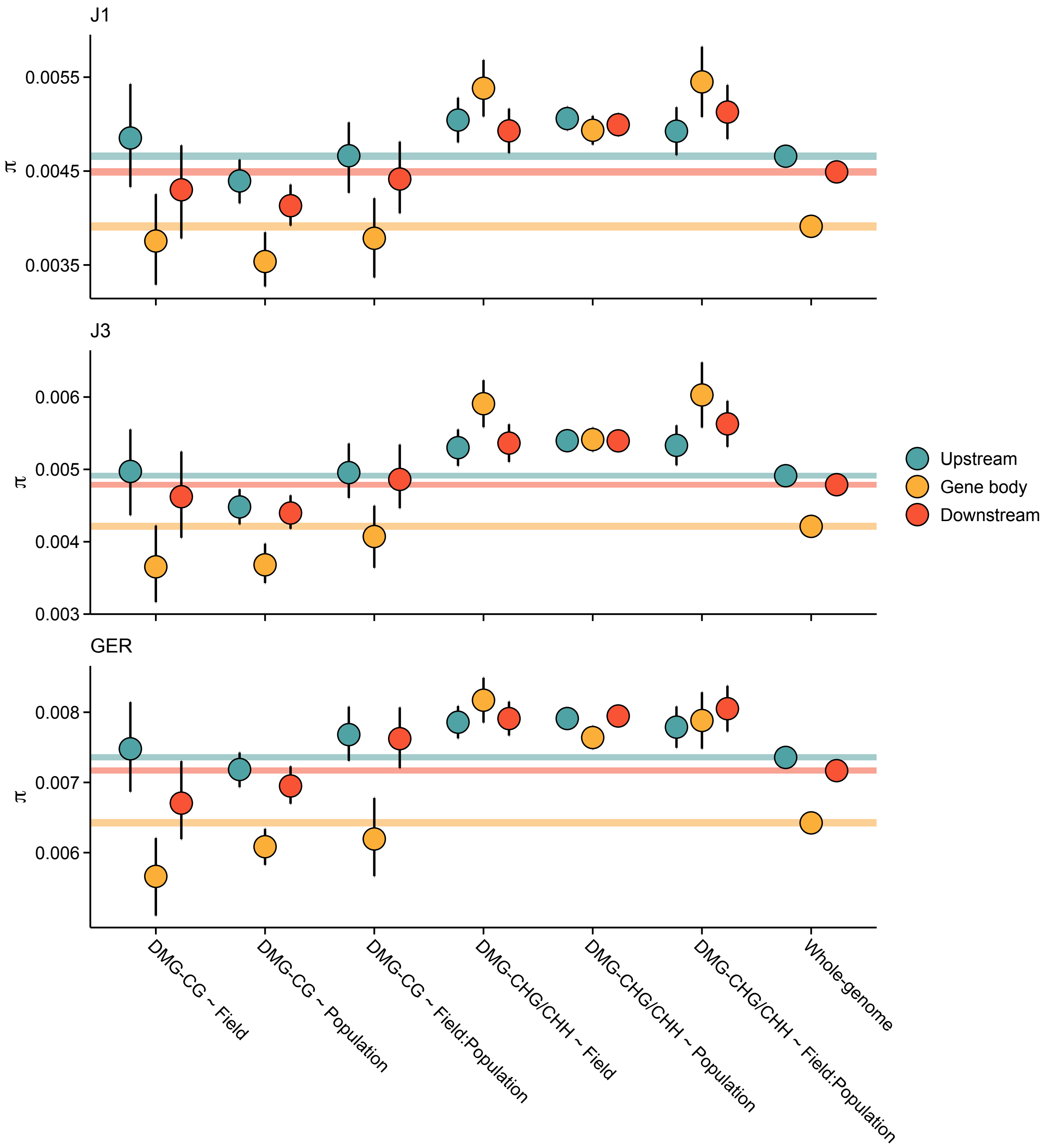


**Figure S10.** Pairwise nucleotide diversity (π) at DMGs. Shown are estimates for gene bodies and 1 kb up- and downstream regions. Error bars show 95% CIs. Shaded areas mark the 95% CIs across all genes.

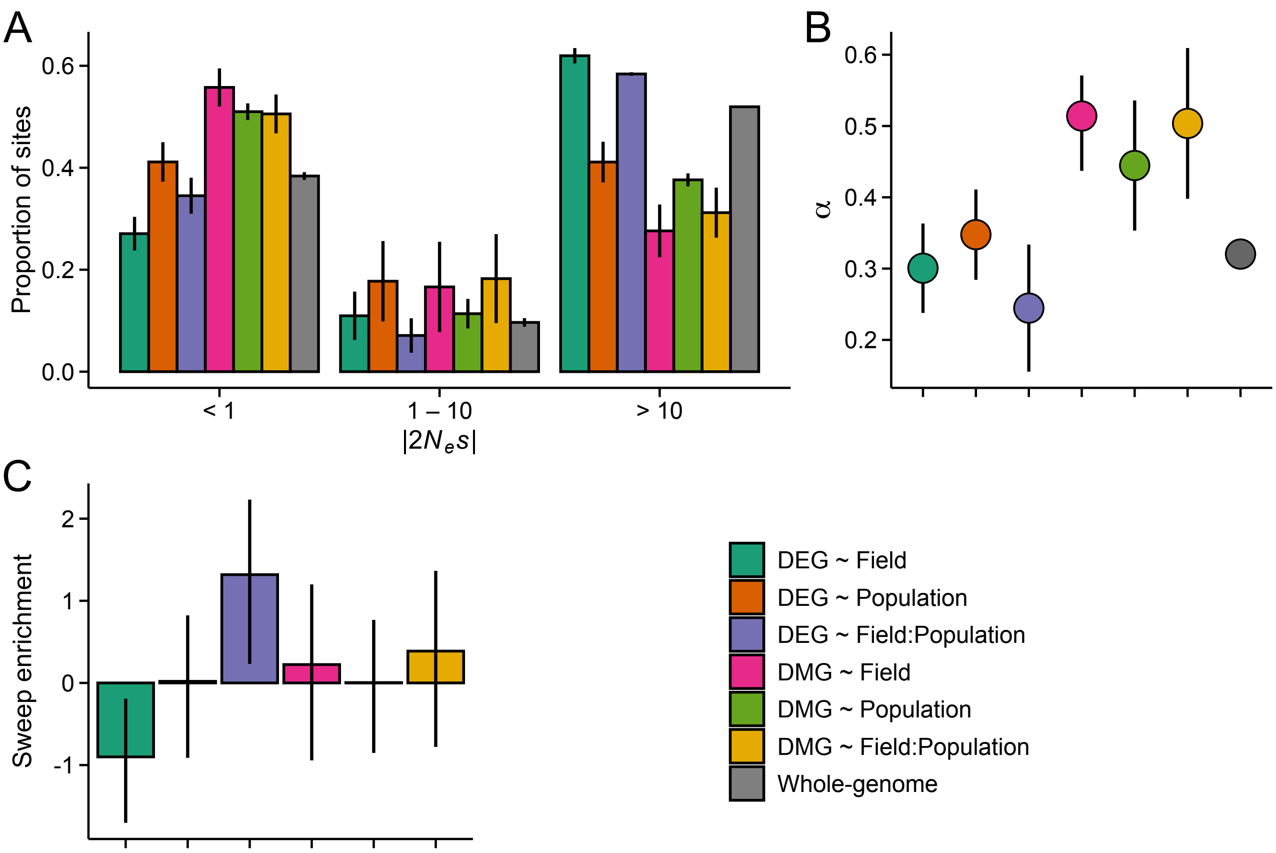


**Figure S11**. The efficacy of negative and positive selection at candidate gene sets. Estimated are shown for J1 ($n$ = 9). **A:** The distribution of fitness effects (DFE) of new nonsynonymous variants. The mutations were divided into three bins based on the strength of purifying selection ($2N_es$): nearly neutral, intermediate, and deleterious, respectively. **B:** The proportion of sites fixed by positive selection ($\alpha$). **C**: The $\log_2$ odds ratio of association between selective sweeps and the candidate gene sets. For all panels, error bars show 95% CIs.

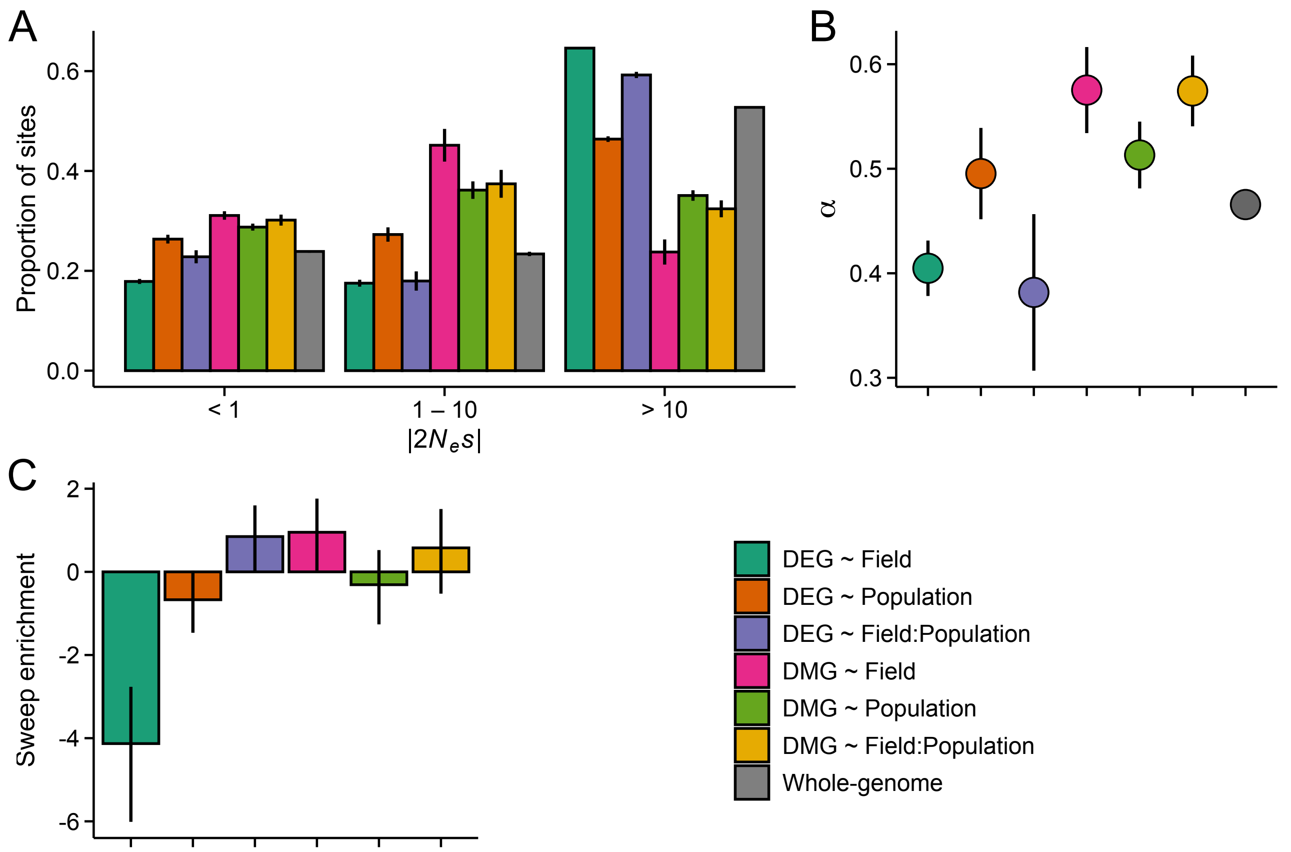


**Figure S12**. The efficacy of negative and positive selection at candidate gene sets. Estimated are shown for GER ($n = 17$). **A:** The distribution of fitness effects (DFE) of new nonsynonymous variants. The mutations were divided into three bins based on the strength of purifying selection ($2N_es$): nearly neutral, intermediate, and deleterious, respectively. **B:** The proportion of sites fixed by positive selection ($\alpha$). **C**: The $\log_2$ odds ratio of association between selective sweeps and the candidate gene sets. For all panels, error bars show 95% CIs.

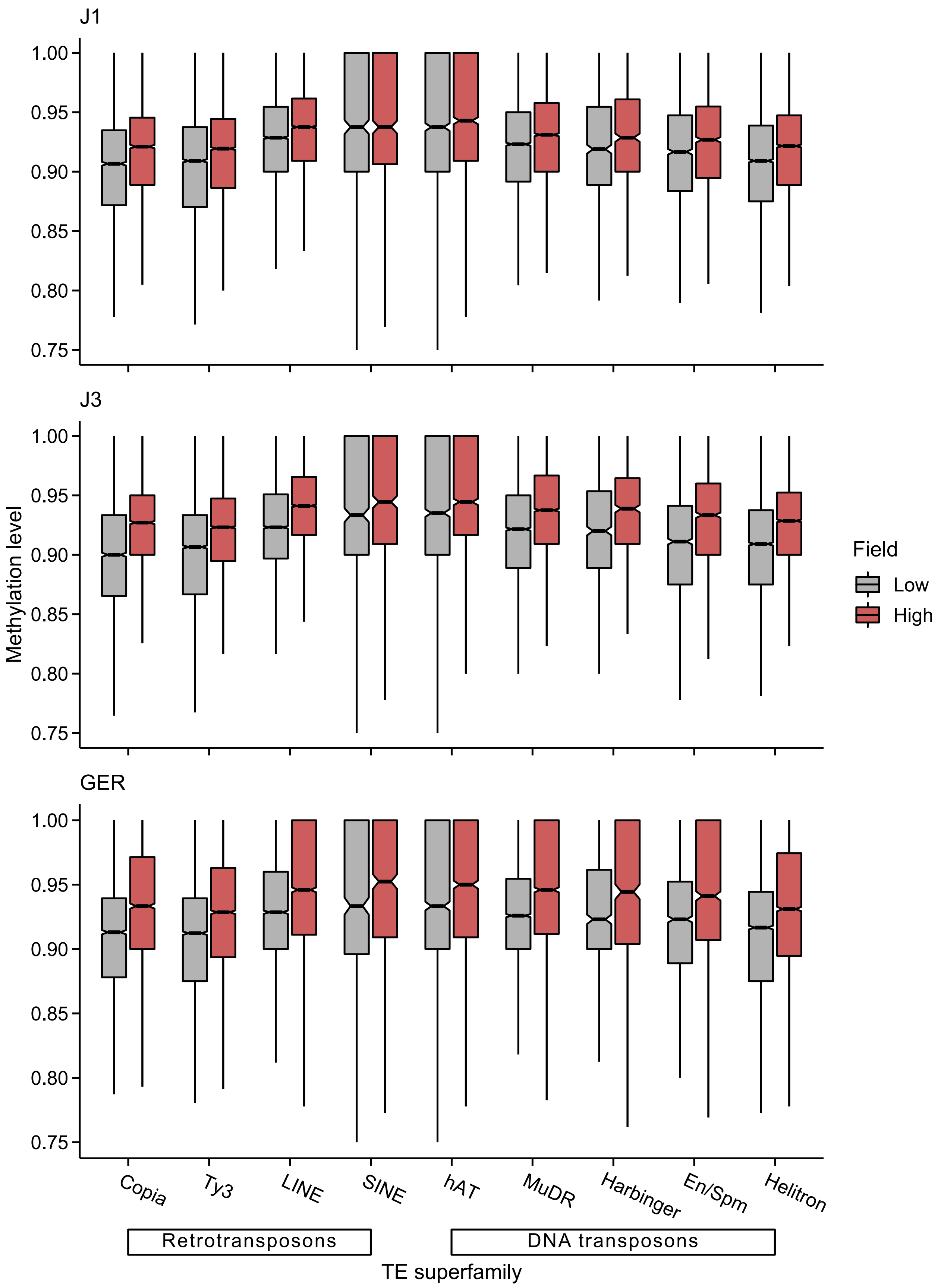


**Figure S13.** CG methylation levels of different TE superfamilies at low- and high-altitude field sites.

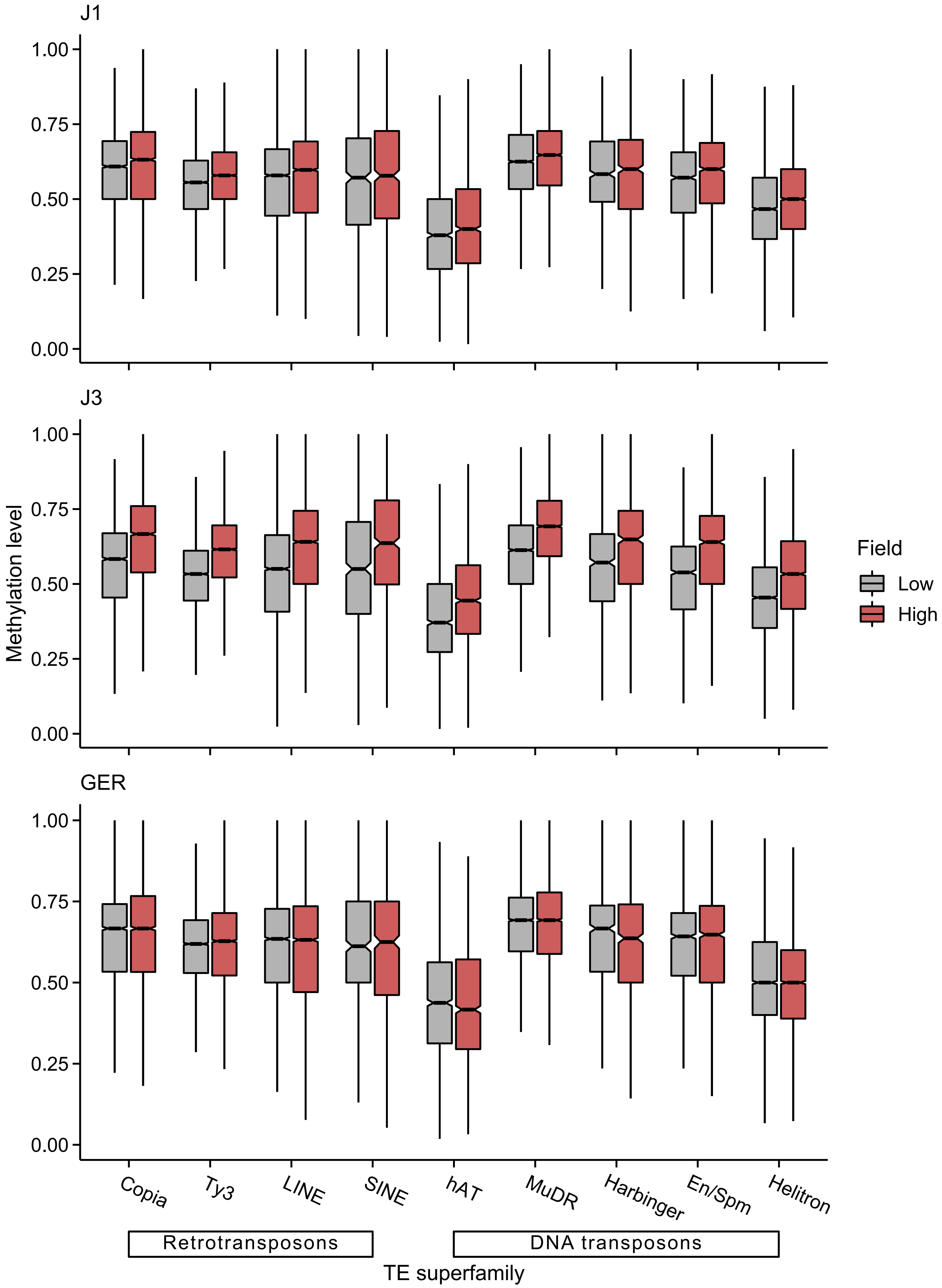


**Figure S14.** CHG methylation levels of different TE superfamilies at low- and high-altitude field sites.

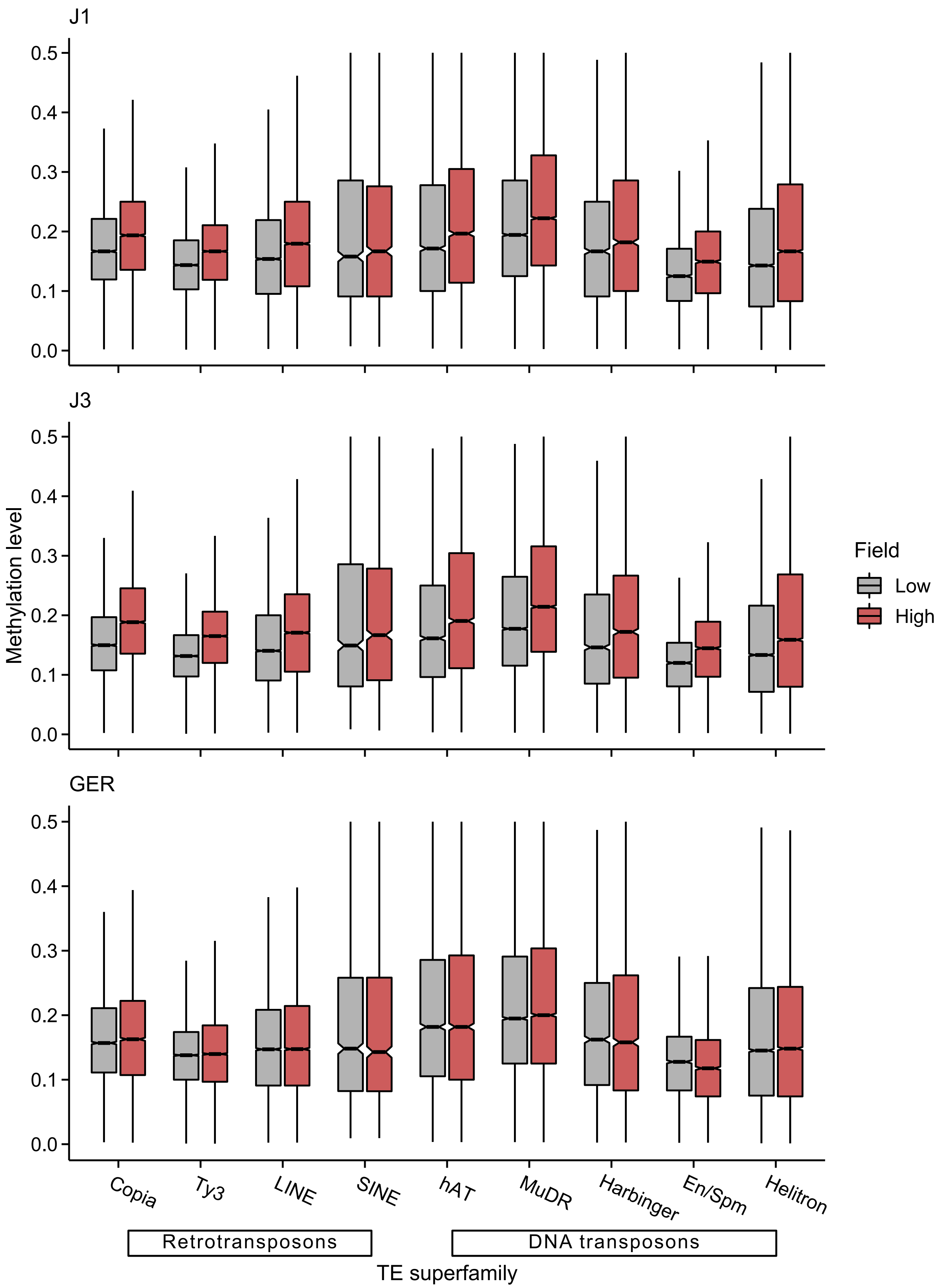


**Figure S15.** CHH methylation levels of different TE superfamilies at low- and high-altitude field sites.

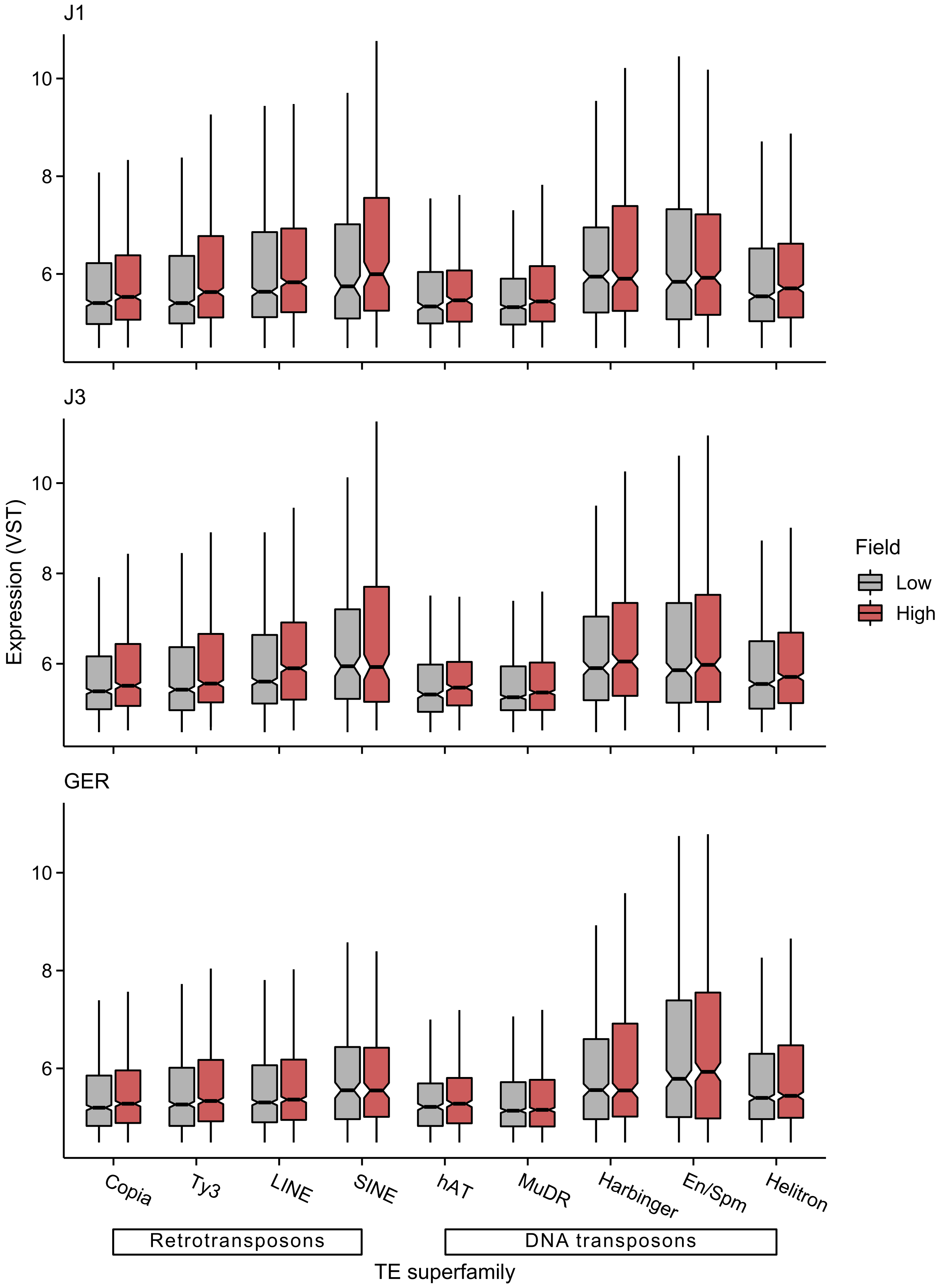


**Figure S16**. Expression levels of different TE superfamilies at low- and high-altitude field sites.

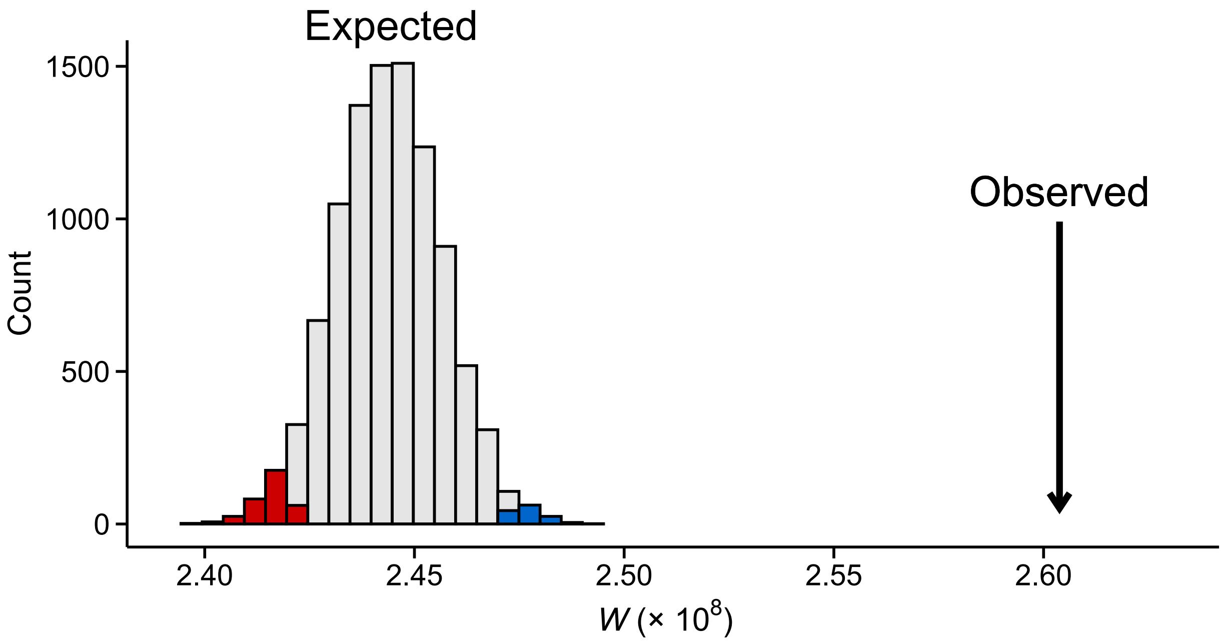


**Figure S17.** Observed TE expression difference between the field sites compared to 10,000 randomly compiled gene sets of equal size. Shown are test statistics (*W*) from Wilcoxon rank-sum tests. Colors indicate $P < 0.05$ (blue = higher expression in the high-altitude field, red = higher expression in low-altitude field). Data were normalized by applying VST on combined gene and TE counts.

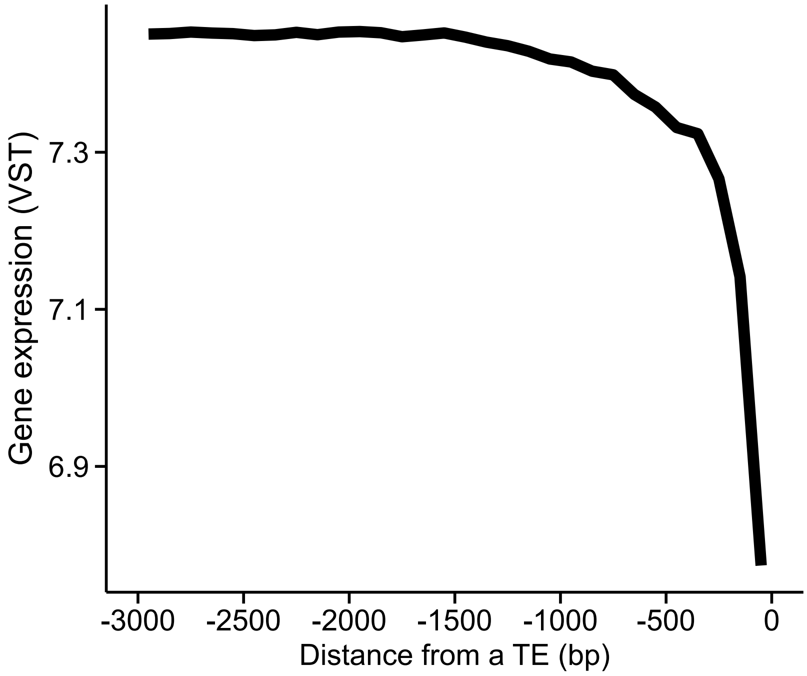


**Figure S18.** The association between gene expression and distance from the closest TE (upstream of TSS). Data were split into 100 bp non-overlapping windows based on their distance and average expression level calculated for each window.

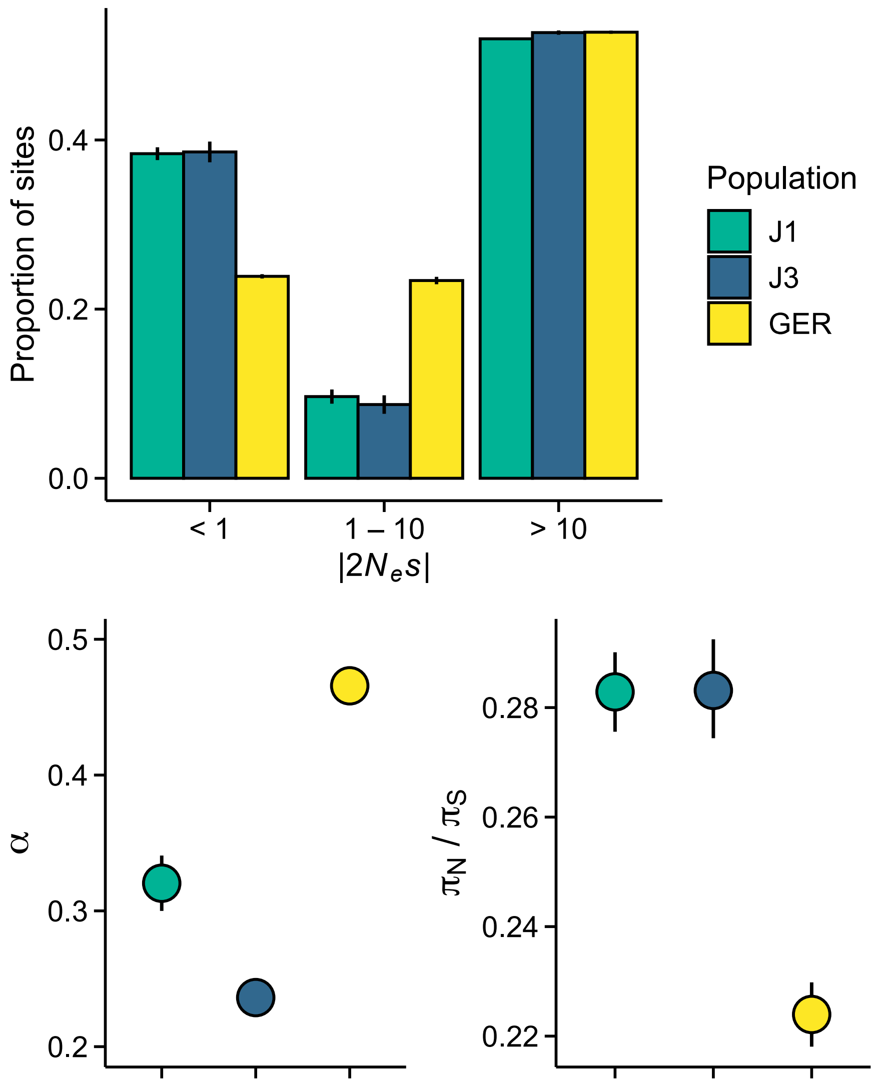


**Figure S19.** Efficacy of selection in the three populations. **A:** The distribution of fitness effects (DFE) of new nonsynonymous variants. The mutations were divided into three bins based on the strength of purifying selection ($2N_es$): nearly neutral, intermediate, and highly deleterious, respectively. **B:** The proportion of sites fixed by positive selection ($\alpha$). **C**: The ratio of nonsynonymous to synonymous nucleotide diversity ($\pi_N/\pi_S$).

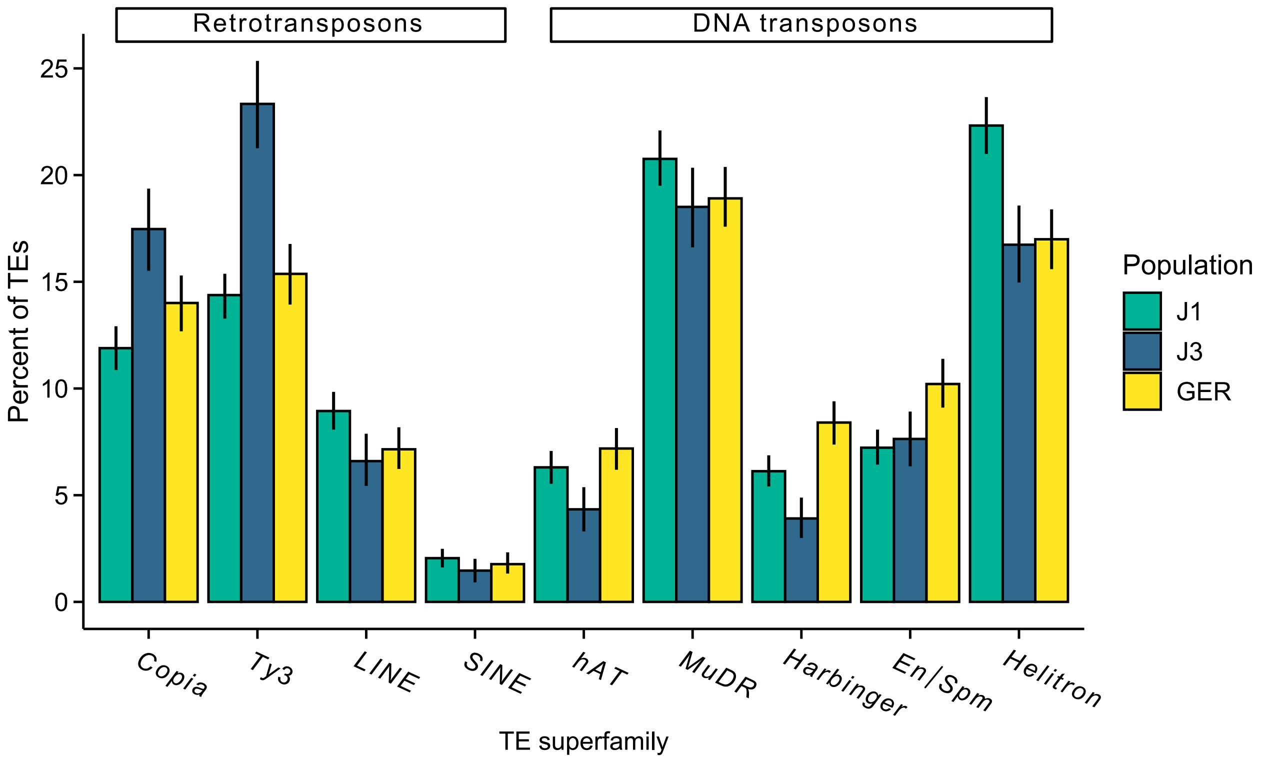


**Figure S20.** The proportion of different TE superfamilies in each population.

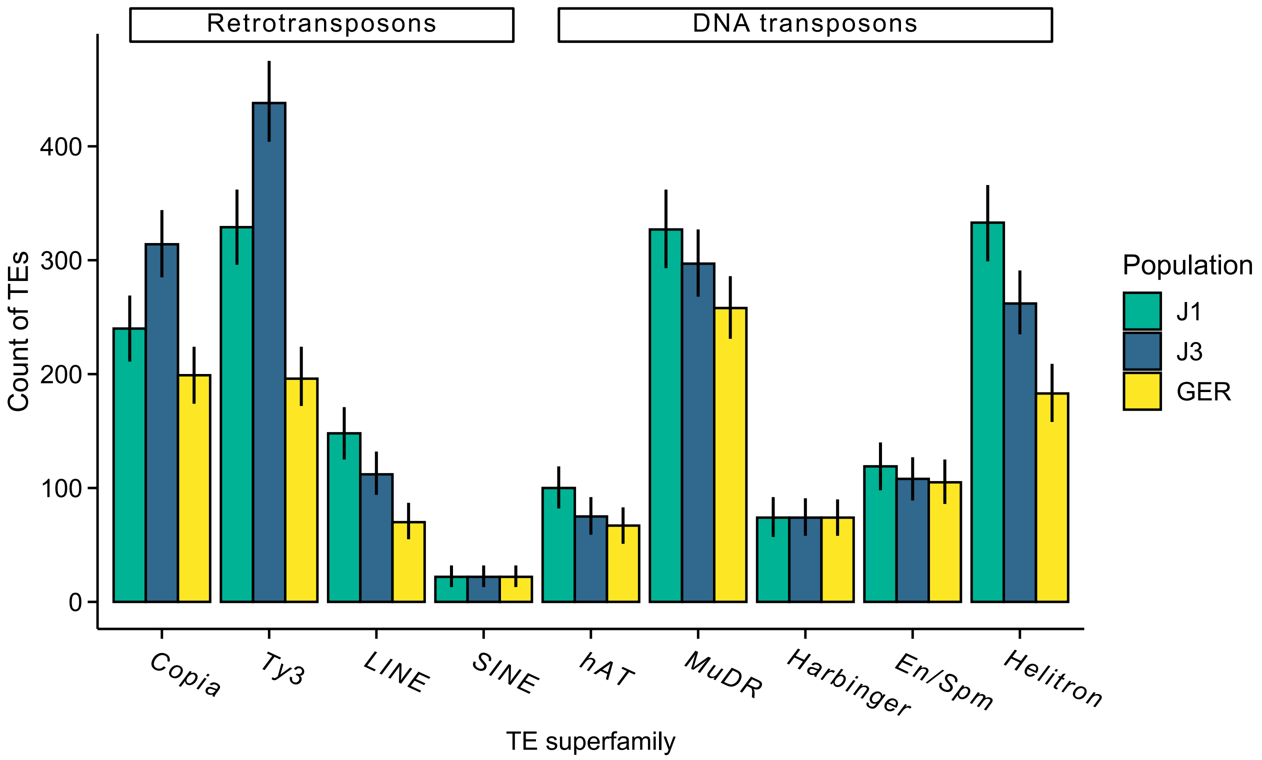


**Figure S21.** The count of different TE superfamilies in each population. The analysis was conducted using the same sample size for each population (*n* = 9) and the same number of aligned read pairs for each sample (40 million).

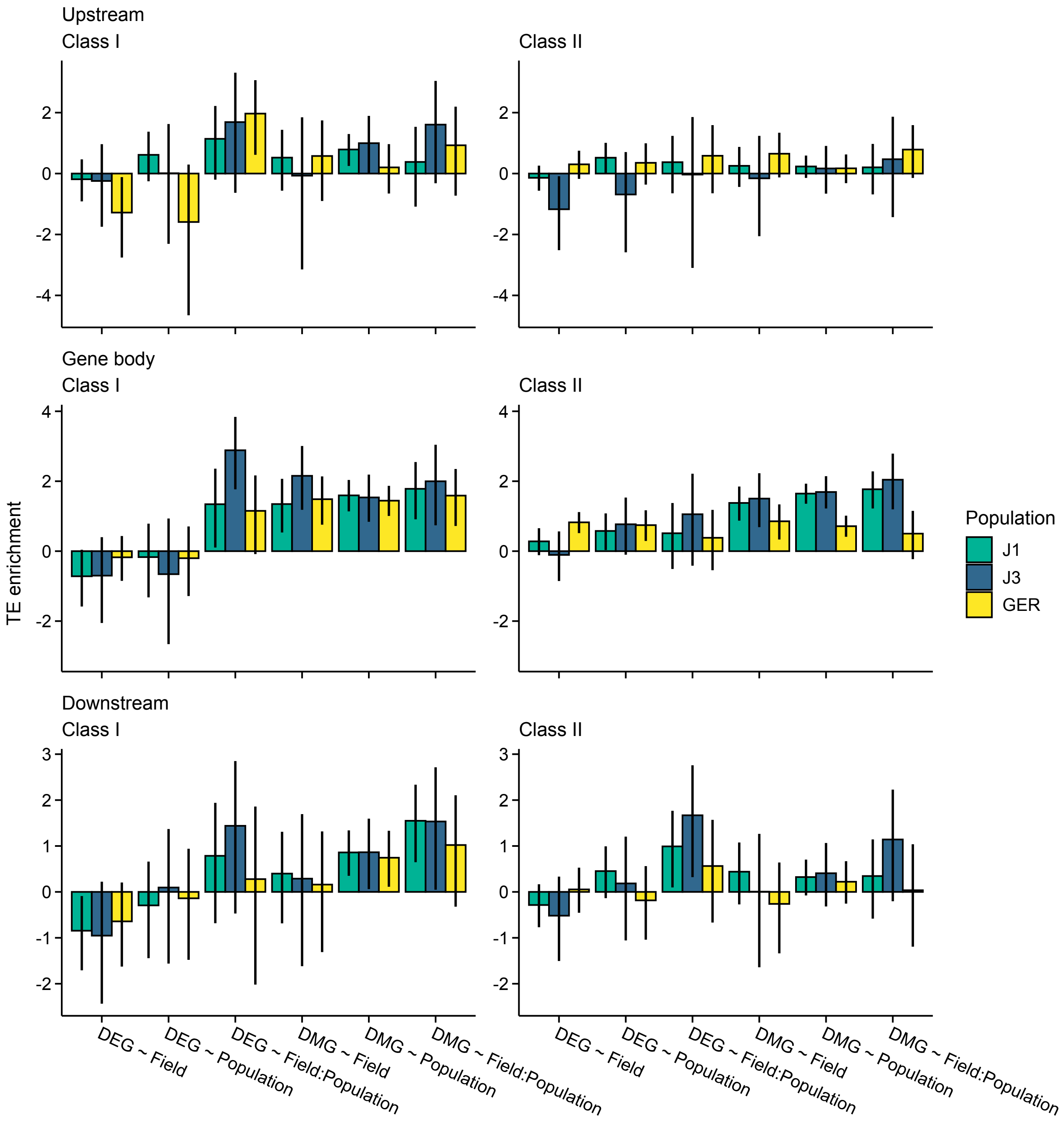


**Figure S22.** The $\log_2$ odds ratio of association between TEs and the candidate gene sets. Shown are estimates for gene bodies and 1 kb up- and downstream regions. Error bars show 95% CIs.

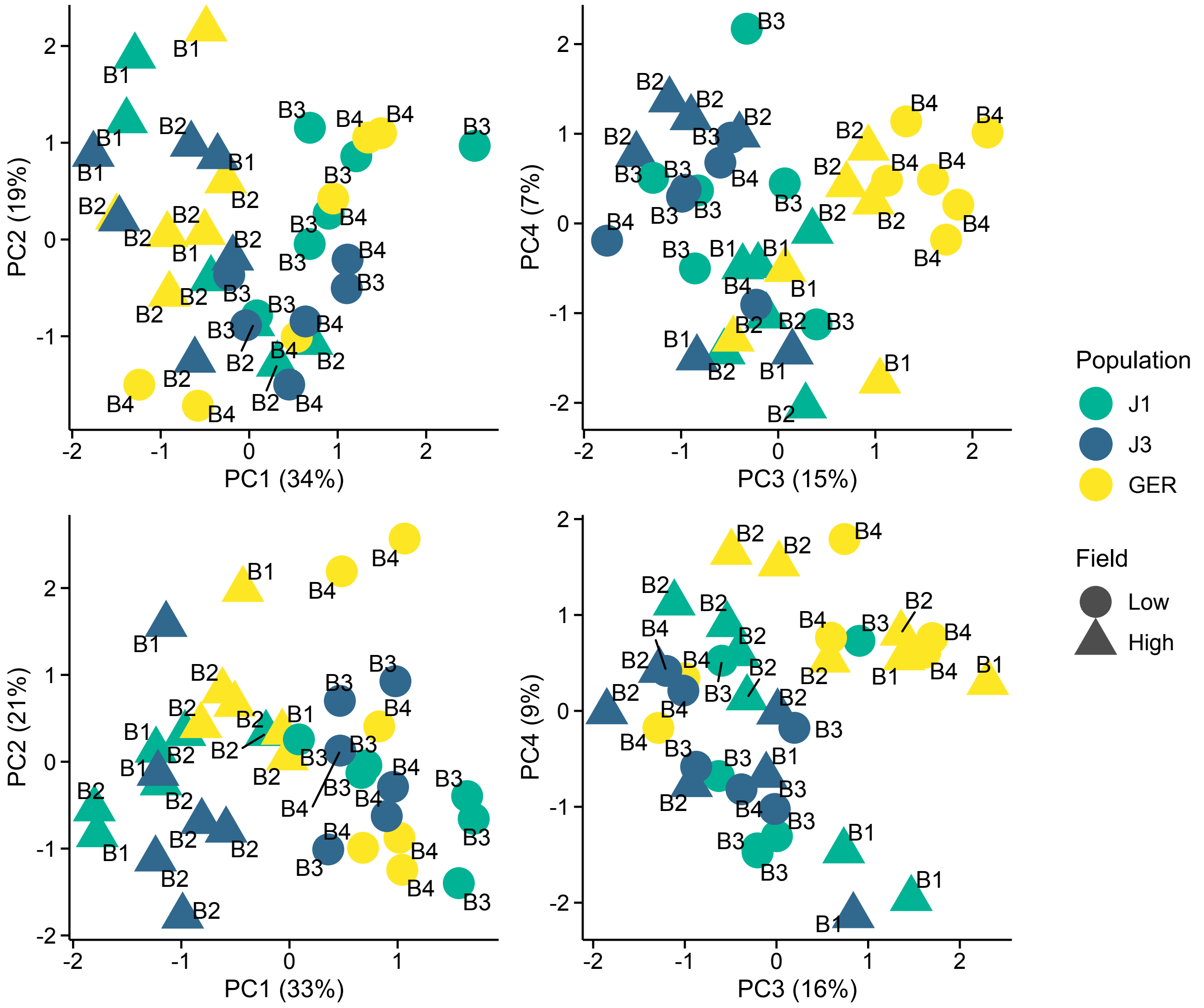


**Figure S23**. Expression variation along first four eigenvectors of a PCA conducted using stably expressed reference genes. The sequencing batch (B1, B2, B3, B4) is marked next to the symbols. Top panels: 20 genes from Czechowski et al. (2005). Bottom panels: 13 genes from Kudo et al. (2016).

**Table S1.** Models for LRTs and the number of identified DEGs and DMGs.

| Gene set | Full and reduced model | Number |
|---|---|---|
| DEG ~ Field | Expression ~ Population + Field<br>Expression ~ Population | 3456 |
| DEG ~ Population | Expression ~ Field + Population<br>Expression ~ Field | 1476 |
| DEG ~ Field:Population | Expression ~ Field + Population + Field:Population<br>Expression ~ Field + Population | 477 |
| DMG-CG ~ Field | CG methylation ~ CH% + Population + Field<br>CG methylation ~ CH% + Population | 112 |
| DMG-CG ~ Population | CG methylation ~ CH% + Field + Population<br>CG methylation ~ CH% + Field | 641 |
| DMG-CG ~ Field:Population | CG methylation ~ CH% + Field + Population + Field:Population<br>CG methylation ~ CH% + Field + Population | 260 |
| DMG-CH ~ Field | CH methylation ~ Population + Field<br>CH methylation ~ Population | 1036 |
| DMG-CH ~ Population | CH methylation ~ Field + Population<br>CH methylation ~ Field | 3580 |
| DMG-CH ~ Field:Population | CH methylation ~ Field + Population + Field:Population<br>CH methylation ~ Field + Population | 680 |

CH = CHG and CHH

CH% = methylation rate at CHG and CHH contexts

**Table S2.** Number of read pairs and mapping rates for each sample.

| ID | Field | Population | Data | Number of read pairs (M) | Mapping rate |
|---|---|---|---|---|---|
| 163 | Low | J1 | RNA-Seq | 61.7 | 0.94 |
| 20 | Low | J1 | RNA-Seq | 24.2 | 0.93 |
| 383 | Low | J1 | RNA-Seq | 14.5 | 0.95 |
| 416 | Low | J1 | RNA-Seq | 22.7 | 0.95 |
| 761 | Low | J1 | RNA-Seq | 21.9 | 0.96 |
| 985 | Low | J1 | RNA-Seq | 36.7 | 0.96 |
| 1601 | Low | J3 | RNA-Seq | 21.1 | 0.95 |
| 1651 | Low | J3 | RNA-Seq | 24.5 | 0.95 |
| 1722 | Low | J3 | RNA-Seq | 31.7 | 0.92 |
| 1727 | Low | J3 | RNA-Seq | 21.8 | 0.95 |
| 238 | Low | J3 | RNA-Seq | 47.6 | 0.96 |
| 750 | Low | J3 | RNA-Seq | 42.9 | 0.95 |
| 2187 | Low | GER | RNA-Seq | 38.4 | 0.94 |
| 2196 | Low | GER | RNA-Seq | 46.7 | 0.90 |
| 2214 | Low | GER | RNA-Seq | 24.4 | 0.93 |
| 2260 | Low | GER | RNA-Seq | 24.1 | 0.91 |
| 2304 | Low | GER | RNA-Seq | 30.4 | 0.94 |
| 2324 | Low | GER | RNA-Seq | 13.9 | 0.92 |
| 1159 | High | J1 | RNA-Seq | 29.2 | 0.96 |
| 1490 | High | J1 | RNA-Seq | 26.6 | 0.95 |
| 1492 | High | J1 | RNA-Seq | 30.7 | 0.94 |

| | | | | | |
|---|---|---|---|---|---|
| 1658 | High | J1 | RNA-Seq | 28.8 | 0.96 |
| 182 | High | J1 | RNA-Seq | 22.9 | 0.95 |
| 2023 | High | J1 | RNA-Seq | 25.4 | 0.94 |
| 1391 | High | J3 | RNA-Seq | 21.3 | 0.95 |
| 1511 | High | J3 | RNA-Seq | 24.4 | 0.88 |
| 1998 | High | J3 | RNA-Seq | 23.3 | 0.94 |
| 608 | High | J3 | RNA-Seq | 20.8 | 0.95 |
| 762 | High | J3 | RNA-Seq | 23.6 | 0.95 |
| 851 | High | J3 | RNA-Seq | 25.8 | 0.90 |
| 2179 | High | GER | RNA-Seq | 25.6 | 0.93 |
| 2274 | High | GER | RNA-Seq | 31.8 | 0.95 |
| 2285 | High | GER | RNA-Seq | 36.0 | 0.94 |
| 2330 | High | GER | RNA-Seq | 17.6 | 0.92 |
| 2432 | High | GER | RNA-Seq | 18.3 | 0.88 |
| 2456 | High | GER | RNA-Seq | 55.3 | 0.93 |
| 163 | Low | J1 | Bisulfite-Seq | 34.4 | 0.62 |
| 383 | Low | J1 | Bisulfite-Seq | 32.8 | 0.67 |
| 761 | Low | J1 | Bisulfite-Seq | 21.3 | 0.65 |
| 985 | Low | J1 | Bisulfite-Seq | 24.7 | 0.63 |
| 1601 | Low | J3 | Bisulfite-Seq | 25.8 | 0.65 |
| 1651 | Low | J3 | Bisulfite-Seq | 46.9 | 0.66 |
| 238 | Low | J3 | Bisulfite-Seq | 27.4 | 0.65 |
| 750 | Low | J3 | Bisulfite-Seq | 25.4 | 0.64 |
| 2187 | Low | GER | Bisulfite-Seq | 34.0 | 0.65 |
| 2214 | Low | GER | Bisulfite-Seq | 30.2 | 0.65 |
| 2304 | Low | GER | Bisulfite-Seq | 73.8 | 0.60 |
| 2324 | Low | GER | Bisulfite-Seq | 26.4 | 0.66 |
| 1490 | High | J1 | Bisulfite-Seq | 33.0 | 0.64 |
| 1658 | High | J1 | Bisulfite-Seq | 36.4 | 0.65 |
| 182 | High | J1 | Bisulfite-Seq | 24.6 | 0.66 |
| 2023 | High | J1 | Bisulfite-Seq | 30.7 | 0.66 |
| 1123 | High | J3 | Bisulfite-Seq | 38.7 | 0.60 |
| 1222 | High | J3 | Bisulfite-Seq | 33.6 | 0.64 |
| 1528 | High | J3 | Bisulfite-Seq | 33.3 | 0.68 |
| 1817 | High | J3 | Bisulfite-Seq | 29.0 | 0.66 |
| 2274 | High | GER | Bisulfite-Seq | 31.9 | 0.60 |
| 2285 | High | GER | Bisulfite-Seq | 33.3 | 0.58 |
| 2297 | High | GER | Bisulfite-Seq | 16.5 | 0.67 |
| 2456 | High | GER | Bisulfite-Seq | 24.4 | 0.64 |

**Table S3.** Bisulfite conversion efficacy for each sample.

| ID | Field | Population | Sequence context | | |
|---|---|---|---|---|---|
| | | | CG | CHG | CHH |
| 163 | Low | J1 | 0.997 | 0.997 | 0.998 |
| 383 | Low | J1 | 0.997 | 0.997 | 0.998 |
| 761 | Low | J1 | 0.994 | 0.997 | 0.998 |
| 985 | Low | J1 | 0.998 | 0.997 | 0.998 |
| 1601 | Low | J3 | 0.991 | 0.995 | 0.997 |
| 1651 | Low | J3 | 0.981 | 0.991 | 0.997 |
| 238 | Low | J3 | 0.997 | 0.997 | 0.997 |
| 750 | Low | J3 | 0.997 | 0.997 | 0.997 |
| 2187 | Low | GER | 0.994 | 0.996 | 0.998 |
| 2214 | Low | GER | 0.998 | 0.998 | 0.998 |
| 2304 | Low | GER | 0.997 | 0.997 | 0.998 |
| 2324 | Low | GER | 0.997 | 0.997 | 0.998 |
| 1490 | High | J1 | 0.986 | 0.990 | 0.995 |
| 1658 | High | J1 | 0.995 | 0.996 | 0.997 |
| 182 | High | J1 | 0.997 | 0.997 | 0.997 |
| 2023 | High | J1 | 0.996 | 0.996 | 0.997 |
| 1123 | High | J3 | 0.968 | 0.980 | 0.995 |
| 1222 | High | J3 | 0.937 | 0.967 | 0.993 |
| 1528 | High | J3 | 0.996 | 0.996 | 0.997 |
| 1817 | High | J3 | 0.995 | 0.996 | 0.997 |
| 2274 | High | GER | 0.995 | 0.996 | 0.997 |
| 2285 | High | GER | 0.994 | 0.995 | 0.997 |
| 2297 | High | GER | 0.991 | 0.993 | 0.997 |
| 2456 | High | GER | 0.998 | 0.997 | 0.998 |

**Table S4.** Species used in estimating nucleotide conservation with GERP++.

| Species | Source |
|---|---|
| *Arabidopsis halleri* | Ensembl Plants |
| *Arabidopsis thaliana* | Ensembl Plants |
| *Arabis alpina* | http://www.arabis-alpina.org/refseq.html |
| *Boechera stricta* | Phytozome |
| *Brassica rapa* | Ensembl Plants |
| *Camelina sativa* | Ensembl Plants |
| *Capsella rubella* | Phytozome |
| *Cardamine hirsuta* | http://chi.mpipz.mpg.de/assembly.html |
| *Carica papaya* | Phytozome |
| *Citrus clementina* | Ensembl Plants |
| *Cucumis sativus* | Ensembl Plants |
| *Eucalyptus grandis* | Ensembl Plants |
| *Eutrema salsugineum* | Phytozome |
| *Fragaria vesca* | Phytozome |
| *Glycine max* | Ensembl Plants |
| *Juglans regia* | Ensembl Plants |
| *Malus domestica* | Ensembl Plants |
| *Manihot esculenta* | Ensembl Plants |
| *Medicago truncatula* | Ensembl Plants |
| *Pistacia vera* | Ensembl Plants |
| *Populus trichocarpa* | Ensembl Plants |
| *Quercus lobata* | Ensembl Plants |
| *Schenkiella parcula* | Phytozome |
| *Theobroma cacao* | Ensembl Plants |
| *Vitis vinifera* | Ensembl Plants |
